# Acoel single-cell transcriptomics: cell-type analysis of a deep branching bilaterian

**DOI:** 10.1101/2020.07.10.196782

**Authors:** Jules Duruz, Cyrielle Kaltenrieder, Peter Ladurner, Rémy Bruggmann, Pedro Martìnez, Simon G. Sprecher

## Abstract

Bilaterian animals display a wide variety of cell types, organized into defined anatomical structures and organ systems, which are mostly absent in pre-bilaterian animals. Xenacoelomorpha are an early-branching bilaterian phylum displaying an apparently relatively simple anatomical organization that have greatly diverged from other bilaterian clades. In this study, we use whole-body single-cell transcriptomics on the acoel *Isodiametra pulchra* to identify and characterize different cell types. Our analysis identifies the existence of ten major cell-type categories in acoels all contributing to main biological functions of the organism: metabolism, locomotion and movements, behavior, defense and development. Interestingly, while most cell clusters express core fate markers shared with other animal clades, we also describe a surprisingly large number of clade-specific marker genes, suggesting the emergence of clade-specific common molecular machineries functioning in distinct cell types. Together, these results provide novel insight into the evolution of bilaterian cell-types and open the door to a better understanding of the origins of the bilaterian body plan and their constitutive cell types.

## Introduction

The emergence and early diversification of bilaterians remains a widely debated subject. Identification and characterization of cell types that makes up animals is key to understanding the diversity of bilaterian tissues and morphologies. The recent advent of single-cell transcriptomics provides a unique technical entry point to view the expression profile of individual cells, enabling us to investigate cell-type identities in various organisms.

Xenacoelomorpha are a recently established phylum of bilaterians (Philippe et al. 2011) whose phylogenetic position has long been a source of debate, because of extreme morphological diversity within the phylum, particularities in their anatomy and the fast-evolutionary rate of their genomes. Some anatomical features of Xenacoelomorpha appear similar to non-bilaterian clades such as the absence of a through-gut, the lack of a coelom, the sole use of cilia for locomotion and the apparent lack of an excretory system. However, they display some core bilaterian features such as a centralized nervous system with diverse levels of organization in Xenacoelomorpha. Mature Xenoturbellida appear to lack a clearly identifiable central nervous system (Raikova et al. 2000) while many members of the subgroup Acoelomorpha contain centralized brains and various amounts of nerve cords (Achatz and Martinez 2012; Martinez, Hartenstein and Sprecher 2017). The variability of tissue architectures has also been demonstrated for other tissues such as the musculature, the mouth and pharynxes and the copulatory organs (Achatz et al. 2013). This apparent flexibility of tissue organization is prominent within the Xenacoelomorpha and highlights the uniqueness of the clade for understanding the mechanisms that regulate the evolution of morphologies and their constitutive building units: the cell types.

Because their anatomical and morphological features seem to share characteristics with those of both cnidarians and bilaterians, this phylum has been at the center of an ongoing debate regarding their use as proxies for an ancestral bilaterian (Baguñà and Riutort 2004; Baguñà et al. 2008; Cannon et al. 2016). Acoelomorphs had been initially thought to be plathelminths due to their similar superficial aspect, but later genetic analysis placed them either as sister group to the remaining bilaterians (Ruiz-Trillo et al. 1999) or within deuterostomes, as a sister group to Ambulacraria (Philippe et al. 2011). A later study taking into account supplementary genomic and transcriptomic data from several species and refined evolutionary models placed Xenacoelomorpha as a sister group to all other Bilaterians (Nephrozoa) making it a candidate phylum to better understand bilaterian origins (Cannon et al. 2016). The use of alternative models of gene evolution have questioned that phylogenetic position (Philippe et al., 2019). In fact, these alternative suggestions of phylogenetic affinities reflect general methodological problems involved in the use of phylogenomic tools and models to reconstruct early diverging clades (Kapli et al. 2020), a problematic that remains unsolved with current approaches.

Few species of acoels (Acoela) have so far been kept in laboratory conditions and used for research. The acoels *Symsagittifera roscoffensis*, *Hofstenia miamia* and *Convolutriloba longifissura* have been used to study photosymbiosis (Dupont et al. 2012; Arboleda et al. 2018), regeneration (Perea-Atienza et al. 2013; Bailly et al. 2014; Srivastava et al. 2014; Sprecher et al. 2015; Srivastava et al. 2017; Gehrke et al. 2019), nervous system morphology and development (Bery et al. 2010; Semmler et al. 2010; Perea-Atienza et al. 2018) and body patterning (Hejnol & Martindale 2008; Hejnol & Martindale 2009; Moreno et al. 2009). A particularly interesting, and tractable, system is the acoel *Isodiametra pulchra.* Since its original description (Smith and Bush 1991), the use of this species in the laboratory has been gaining acceptance because of their easy maintenance and of the availability of different technologies, from in situ and immunochemistry to the gene knockdown using RNAi interference (DeMulder et al. 2009; Moreno et al. 2010) and transcriptome sequences (Cannon et al. 2016; Brauchle et al. 2018). *Isodiametra pulchra* has been instrumental in developing many key studies of the Xenacoelomorpha, for instance: detailed descriptions of stem-like cells (De Mulder et al. 2009), the nervous system (Achatz & Martinez 2012), mesoderm (Ladurner et al. 2000; Rieger et al. 2003; Chiodin et al. 2013) or excretory cells (Andrikou et al. 2019).

Recent advances in single-cell RNA sequencing technologies have enabled the thorough description of the full repertoire of cell types (cell atlas) of various organisms by defining cell types as groups of cells clustered based on their RNA expression profiles. This has been done in many animals including both non-bilaterians (Sebe-Pedros et al., 2018a, Sebe-Pedros et al., 2018b, Siebert et al., 2019) and some bilaterian “model” organisms, for instance: the planarian *Schmidtea mediterranea* (Fincher et al. 2018; Plass et al. 2018; Swapna et al. 2018), *Drosophila melanogaster* (Karaiskos et al. 2017), *Mus musculus* (Han et al. 2018), the nematode *C. elegans* (Packer et al. 2019) and the annelid *Platynereis dumerilii* (Achim et al. 2018) among others. These different studies have all revealed a surprising level of cell type heterogeneity and the presence of previously unknown cell-types in these animals.

In this study, we deep-sequenced whole *Isodiametra pulchra* hatchlings at a single-cell resolution to reveal the diversity of cell types in this representative of the enigmatic phylum Xenacoelomorpha with the aim of understanding how these different cells contribute to the organization of its specific body plan. We find a rich diversity of cell types corresponding to well-known bilaterian tissues. Particularly remarkable is the diversity within the nervous system of *I. pulchra*. We further describe cells involved in diverse metabolic activities such as digestion and excretion. Interestingly, we find a variety of putative secretory cells that may play a role in defense and innate immunity and others that are putatively involved in the secretion of adhesive substances. Interestingly, while most cell types of *Isodiametra pulchra* express well known cell-type markers shared with other animal clades we also observe large numbers of co-expressed genes, which appear only to be present in Xenacoelomorpha (*Symsagittifera roscoffensis* and *Xenoturbella bocki*) suggesting the presence of a phylum-specific group of genes contributing to the establishment of cell-type identities among the Xenacoelomorpha.

## Results

### *Isodiametra pulchra* single-cell transcriptomes depict a repertoire of 10 major cell type categories

The analysis of single-cell RNA sequencing of whole *I. pulchra* hatchlings resulted in an estimate of 14,864 recovered cells, after aggregation of two independent experiments, resulting in approximately a 14x coverage of the expected cell number in a whole hatchling that we estimated to be in the range of 900-1000cells. This latter number was determined by automated counting of stained nuclei in a 3D-reconstructed confocal microscopy stack of a whole hatchling (Fig 1C). The median Unique Molecular Identifiers (UMIs) and gene-per-cell estimates were of 608 and 405 respectively with a total of 21,597 genes detected. The UMI values are consistent with those of other studies, with the understanding that these numbers are extremely variable from one species to the other and sometimes even from one experimental condition to another in the same species (Fincher et al. 2018; Sebe-Pedros et al. 2018a, 2018b; Swapna et al. 2018). The mean reads per cell was of 49,404; post-normalization. The data was filtered to only include cells with a gene-per-cell count of 200 to 2000 to exclude cells of poor quality and possible multiplets (Supp data, S2). Cells were clustered using 25 principal components selected in base of the assessment of an elbow plot that ranks principle components according to the percentage of variance that they explain (Satija et al. 2018; Supp. data S2) and with a resolution of 2.5. This resulted in the detection of a total of 42 clusters (Supp. data S2). Identity was assigned to these clusters by analyzing the best markers of each cluster, which are the genes that were most differentially upregulated in one specific cluster with respect to all others.

**Figure 1:**
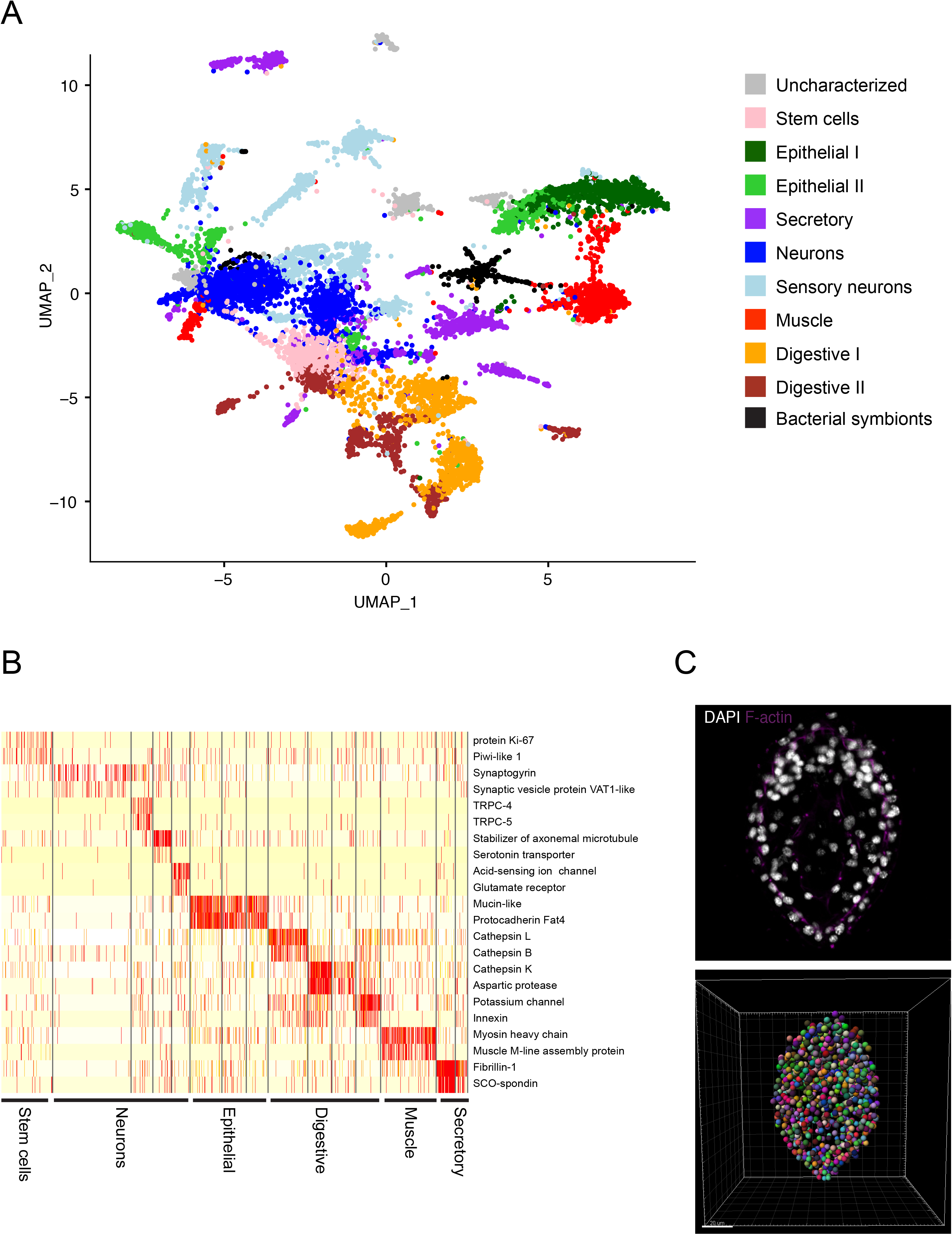
Clustering of *Isodiametra pulchra* cells depicts a repertoire of 10 major cell type categories. (A): UMAP of all cells showing the assignment of all 42 cell clusters to cell-type categories (B): Heatmap showing cell-type specific markers for some of the cell categories (C): Confocal image showing staining of nuclei in an *Isodiametra pulchra* hatchling and corresponding visualization of automated nuclei counting in a three dimensional reconstruction.

Clusters were manually annotated and fitted into ten subjectively defined categories based on the predicted function of identified markers (Fig 1A) which also included a category of uncharacterized cell types. In addition we identified what appeared to be prokaryotic sources, based on the identification of some bacterial rRNA sequences. These prokaryotic sequences could reflect the presence of endosymbionts in *Isodiametra pulchra*, since the same transcripts were found in both the single-cell pools and the RNA sequences used for transcriptome assembly. A few representative markers of each category were plotted to visually assess their enrichment within each cell cluster (Fig 1B). Defined categories include stem-cells, neurons, two distinct types of digestive cells, two distinct types of epithelial cells, secretory cells and muscle cells.

### Stem cells express piwi-like genes and conserved proliferation markers

An interesting problematic in the field of stem cell biology is the putative convergence of neoblast phenotypes in the Platyhelminthes and the Acoela. Here, we adress this topic by analyzing piwi positive cells (a specific marker for stem cells) in *Isodiametra pulchra*. Specifically, a population of cells with stem cell-like profiles was identified based on the broad expression of *piwi-like 1* and *piwi-like 2* (De Mulder et al. 2009); those profiles are mainly aggregated in two clusters of cells (Fig 2A, 2B) both of which express a variety of other genes involved in proliferation, cell growth, DNA replication, organelle biosynthesis and protein synthesis (Fig 2A). While most of these markers are concentrated within two clusters many other cell types seem to have cells sharing an elevated expression of piwi-like genes (Fig 2B).

**Figure 2:**
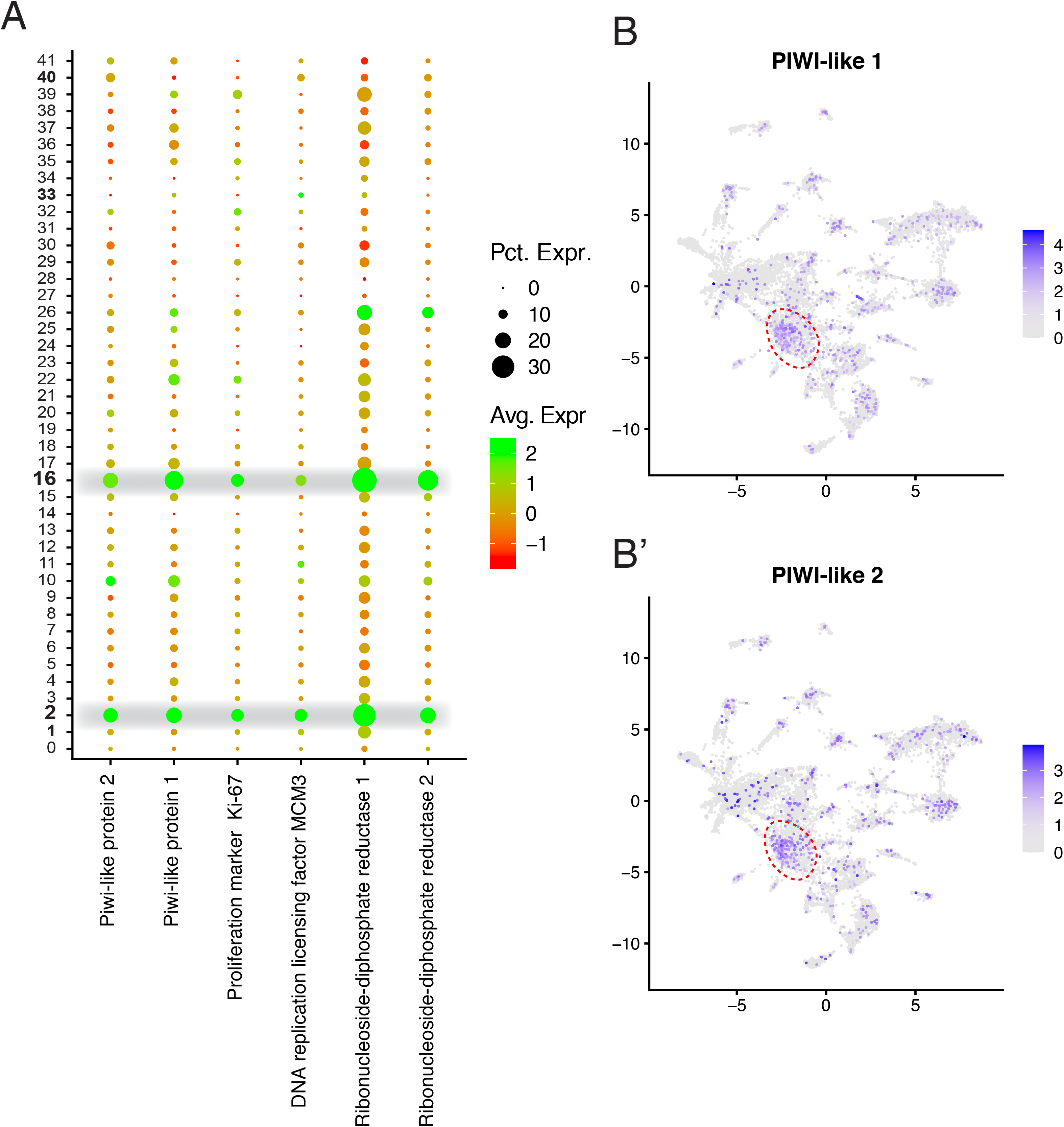
Stem cells express piwi-like genes and conserved proliferation markers. (A) Dotplot showing expression of Stem-cell and proliferation markers across all 42 cell clusters. (B) Feature plots showing the expression of PIWI-like 1 and PIWI-like 2 across all cells. Red dotted line highlights clusters 2 and 16.

### *Isodiametra pulchra* nervous system displays a high diversity of sensory cell types

Because of the high number of clusters with neuronal profiles (10 clusters in total), all the cells from these clusters were batched together and sub-clustered using 10 principal components and a resolution of 0.6 to obtain finer distinctions between neuronal subtypes. This procedure resulted in a total of 12 neuronal sub-clusters. The clusters were sorted into categories defined based on the expression of certain specific markers (Fig 3A). A large population of cholinergic neurons was identified based on the broad expression of *choline acetyl-transferase* (Slemmon et al. 1991; Kim et al. 2006; Achatz and Martinez 2012). Presumed neuronal precursors or differentiating, immature neurons were characterized based on the expression of growth-factor-related genes (*epidermal growth factor like-1, cd63)*, DNA synthesis (*elongation factor 1A2*, *DNA primase/helicase*) and mitosis markers (*microtubule-associated proteins RP/EB*). Another cluster was characterized by the expression of several Transient Channel Potential (TRPs) orthologs (*trpc5a, trpc5b, trpc4, trpa1*), which are classically associated with sensory functions (Rubin 1989; Brauchi et al. 2006; Montell & Peng et al. 2015; Kozma et al. 2018). However, due to the absence of other clear sensory markers in these clusters, they were simply called TRP^+^ neurons. To visualize the domains of expression of TRPC5 the expression was assessed using a single-molecule fluorescent In Situ hybridization (smFISH) technique (Stellaris, LGC biosearch technologies). We observed a clear expression domain located in the anterior tip of the animal (Fig 3C) and in the periphery of the brain. The revealed pattern indicates that some TRPC5^+^ cells are closely connected to the brain, either as a sensory input or as part of the CNS itself. The absence of photosensory neurons in this species has been mentioned by different authors but has never been studied in detail. In this context, we performed an additional analysis of TRP^+^ neurons with the aim of looking for possible correlations between TRP and Opsin expression levels (Supp fig S3). This analysis revealed that TRP^+^ neurons also express more opsins than any other clusters suggesting that some of these cells could indeed be functioning as photoreceptor neurons.

**Figure 3:**
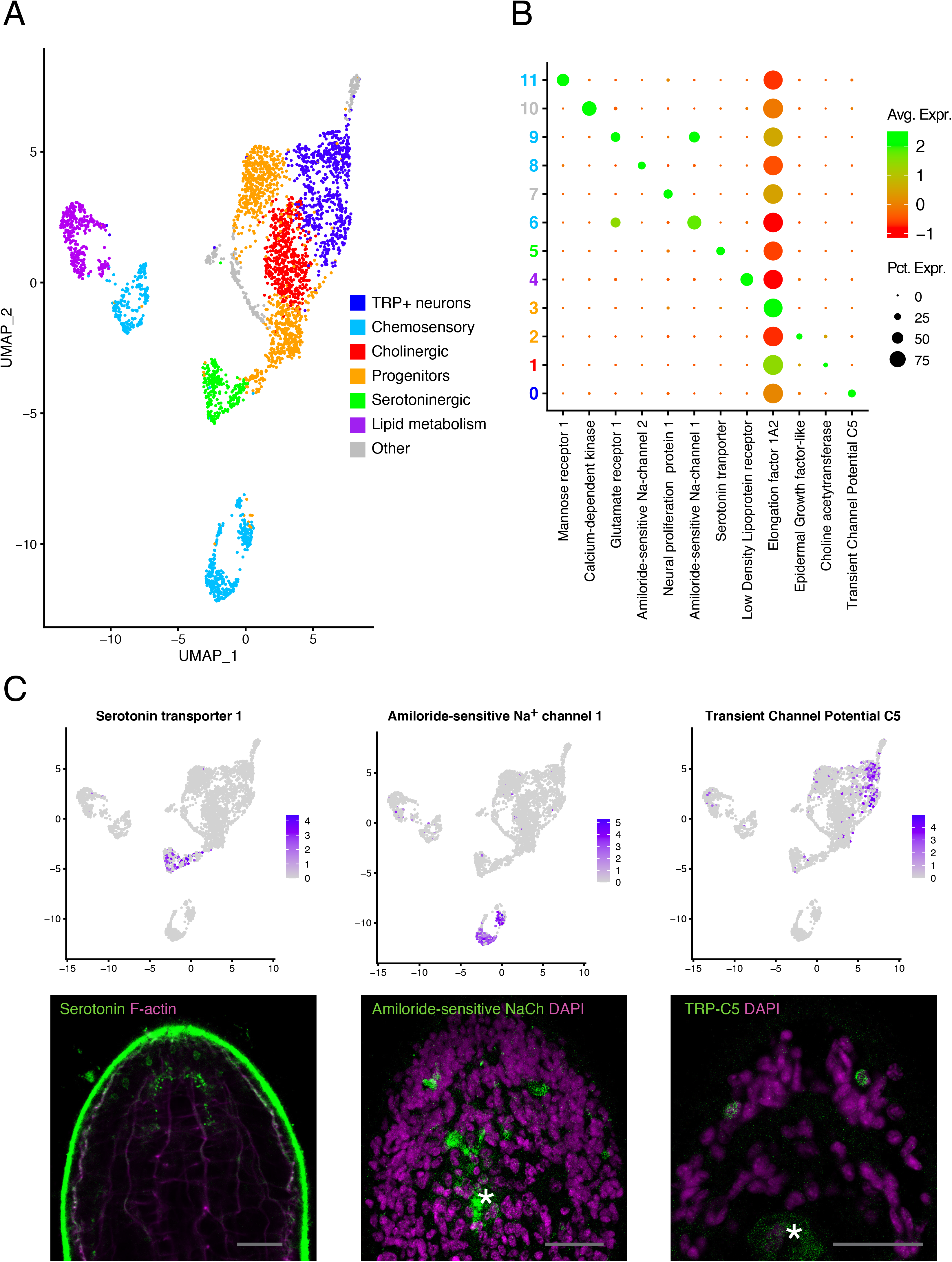
Isodiametra nervous system displays a high diversity of sensory cell types. (A) UMAP showing neuronal sub-clusters and their assigned categories (B) Dotplot showing the expression of main markers of each of the 12 clusters. Cluster numbers are colored according to their assigned categories. (C) Featureplots showing the expression of different specific neuronal markers with corresponding images showing immunostaining with anti-serotonin antibody and smFISH against Amiloride-sensitive Na+ channel 1 and TRP-C5. Scale bars = 25 μM.

Serotoninergic neurons were identified by the expression of a serotonin transporter (*sc6a4*) (Blakely et al. 1991; Corey et al. 1994; Chang et al. 1996). High expression of the microtubule stabilizer *saxo2* indicated that these cells are likely ciliated, which could serve a mechanosensory function. Immunostainings with antibodies against serotonin highlighted a population of serotoninergic neurons in the brain of the animal with cell bodies located towards the anterior tip (Fig 3C). These neurons display a bipolar morphology that appears to connect the tip of the animal to the CNS supporting their presumed function as sensory cells. Immunoreactivity of serotoninergic neurons have been documented in previous studies (Achatz and Martinez 2012; Dittman et al. 2018) but were rather described as a component of the CNS and not necessarily as sensory cells.

Four clusters of distinct cell populations were identified as chemosensory, based on the expression of different combinations of amiloride-sensitive sodium channels and acid-sensing sodium channels (Supp. table 2). Chemosensory cells could be resolved into two distinct populations with one expressing predominantly glutamate receptors (*NMDA-1, Glr1, Grl2*) and the other expressing predominantly acetylcholine receptors (*AChR-1, AChR-2, AChR-3*, Fig 3A). This suggests that these sensory cells likely do not only have the ability to respond to chemical stimuli from the environment but also to be modulated by other neurons. The detection with smFISH for the amiloride-sensitive sodium channel *scnng* revealed instances of expression in the proximity of the brain but also in close proximity to the mouth opening (with often observed background in the digestive syncytium) (Fig 3C). The expression pattern is consistent with the presumed function of these cells to sense chemical compounds in the environment during navigation but also when grazing on algae.

One cluster of identified cells is possibly involved in providing nutrients to the nervous system since they express several lipoproteins receptors (*ldlr1, lrp2*). The function of these cells is uncertain, but they could be providing metabolic support to the nervous system in a glial cell-like manner but due to the broad expression of many lipoprotein receptors in other cell types it is not sufficient to support that hypothesis. However, since tentative glial cells have been identified already in another acoel, *Symsagittifera roscoffensis* (Bery et al. 2010), this remains a plausible hypothesis.

### Digestion and nutrient transport can be shared within a single cell type

All clusters previously identified as digestive cells (I & II) were analyzed. Digestive cells were characterized by the expression of various digestive enzymes such as peptidases and lipases (Fig 4A) and interestingly by the expression of a broad variety of cathepsins known for their catabolic activity but usually inside lysosomes. Cathepsins have been previously described as markers for a novel cell type in *Schmidtea mediterranea* (Fincher et al. 2018; Swapna et al. 2018) however, we show here with the co-expression of cathepsins and other digestive enzymes that their function in acoels is likely to contribute to digestion of food. Whether this digestion happens through secretion of these enzymes into the digestive syncytium or intracellularly remains unknown. In the case of *Isodiametra pulchra* as well as many other acoelomorphs, digestion is supposed to be carried out by the digestive syncytium, a very large polynucleated cell capable of engulfing and digesting food (Gavilán et al. 2019); though it is unclear whether additional cells in the periphery are also involved. Our experimental procedure was designed to dissociate and isolate individual cells and therefore excluded the digestive syncytium from the experimental system. Strikingly, our data shows clearly that there are other cell types also contributing to the digestion of nutrients. These cells are likely to be layering the digestive syncytium to either take in food particles from the syncytium and digest them intracellularly and/or directly secrete digestive enzymes into the syncytium. This assertion is further supported by the expression pattern of cathepsin B shown by In Situ hybdridization in cells surrounding the mouth opening (Fig 4C).

**Figure 4:**
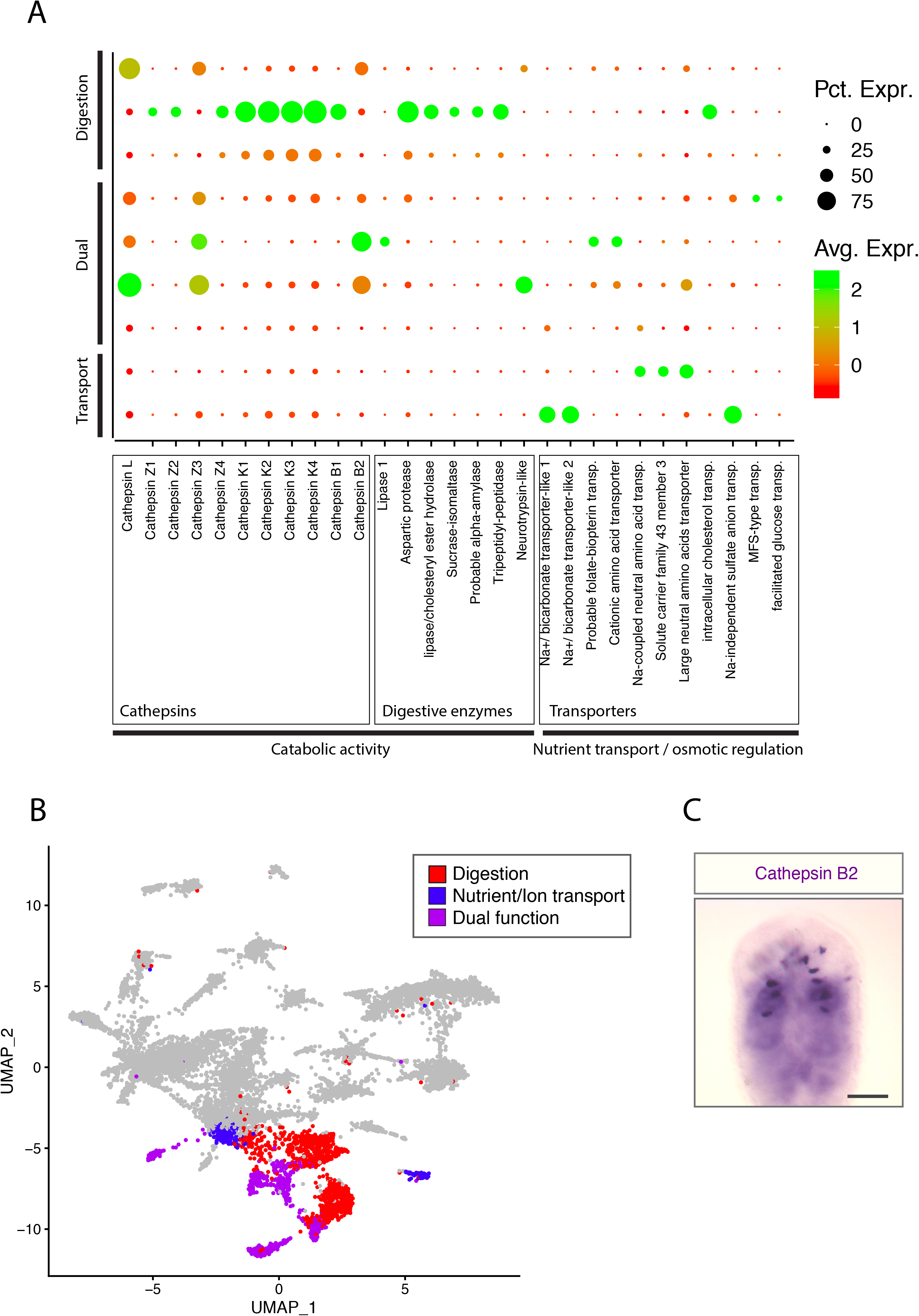
Digestion and nutrient transport can be shared within a single cell type. (A) DotPlot showing the expression of specific markers for 9 identified as “digestive I”, “and “Digestive II”. Genes are sorted by functional categories. (B) UMAP highlighting the cell types involved in digestion, nutrient transport or both (C) ISH showing the expression pattern of cathepsin B-2. Scale bar = 50μM.

Cells expressing nutrient and ion transporters were identified (Fig 4A). These cells could be serving the function of both distributing nutrients to other cells and/or, like an excretory system, to filtrate and reabsorb necessary elements while discarding waste. The existence of such an excretory cell type in *I. pulchra* was previously proposed in Andrikou et al. (2019). We looked for the genes tested as excretory markers in that study and found high expression levels of *nephrin/kirre* and *aquaporin b* in a large population of these transporter-rich cells, consistent with their hypothesis.

Several clusters express a mix of digestive enzymes and transporters suggesting an ability to serve both functions of secreting digestive enzymes and taking up the processed nutrients suggesting the presence of cellular variegated phenotypes (Fig 4A).

### Acoel epithelial and secretory cells predict high functional diversity

Epithelial and secretory cells were analyzed together because of the shared expression of some common markers (spondins, lectins, cadherins, fibrillins), even though we have seen that they are highly diversified and composed of many subtypes. These populations of cells were separated into three categories: epithelial, secretory and motor ciliated cells, each containing several sub-categories based on differential marker expression (Fig 5A, 5B). Epithelial cells were mainly defined by the expression of the transcript for *mucin-like*, a secreted protein characteristic of epithelia in many animal species (Marin et al. 2008), and of multiple cadherins and protocadherins responsible for cell-cell adhesion, critical for epithelium formation. These cells also expressed *sortilin-like* receptors (Mazzella et al. 2019), which are broadly studied in vertebrates but are of unknown function in invertebrates. Interestingly, this cell type category expresses a *myosin-11* ortholog that could be an indicator of cell contractility.

**Figure 5:**
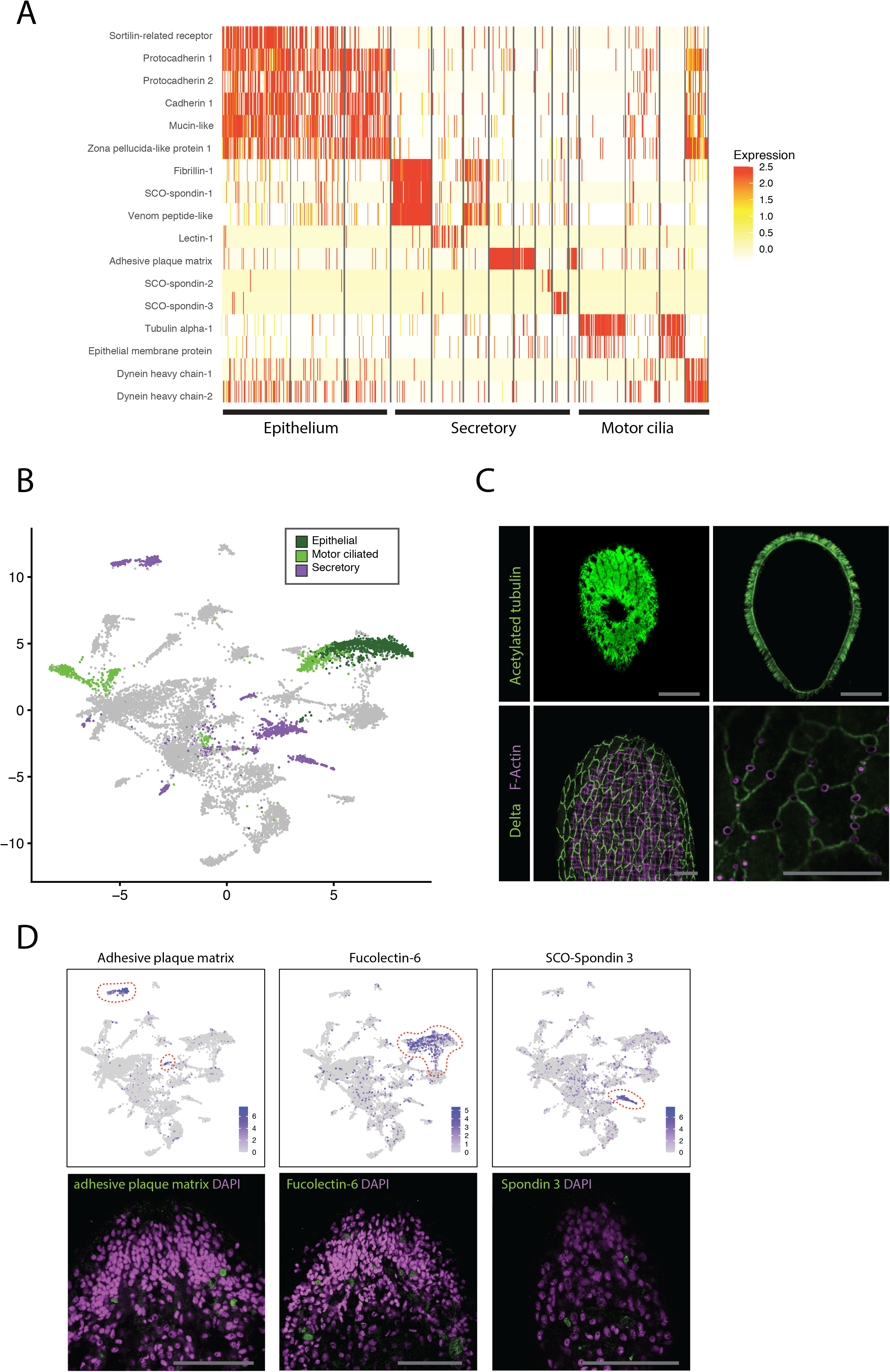
Acoel epithelial and secretory cells predict high functional diversity. (A) Heatmap showing the expression level specific markers for the 15 cell clusters identified as “Epithelial I”, “Epithelial II” and “Secretory”. Rows represent genes and columns represent cells. (B) UMAP highlighting the cell types presumed to be epithelial, motor ciliated and or secretory. (C) Immunostainings using anti-acteylated tubulin antibodies and anti-dDelta antibodies that highlight the heavy ciliation of the animal and the outline of the *I. pulchra* epithelial cells respectively. Scale bars = 25μM (D) Feature plots showing the expression of specific markers for secretory and epithelial cells across all cells and their corresponding smFISH images. Scale bar = 50μM.

Secretory cells were one of the most diverse group of identified cell types in our dataset. Many genes identified in this clusters had no known orthologues making it difficult to assess cell-type identity, but they nevertheless shared some conserved core markers: most of these cells express at least one type of fibrillin, and/or spondin. Expression of lectins, fucolectins, as well as cysteine-rich venom-related proteins (*va5, vpl1*), suggest a possible involvement in defense against predators and/or pathogens. A subset of these cells strongly expressed an ortholog of an *adhesive plaque matrix protein,* a protein known to form a strong glue-like substance that is molded into holdfast threads in the mussel *Mytilus galloprovincialis* (Inoue & Odo 1994).

Motor ciliated cells were characterized by the very high expression of tubulins, presumably involved in motile cilia formation as well as dyneins known to be involved in the movements of those cilia. Dyneins are also frequently found in other epithelial cells (Fig 5A), indicating that they might not be the only cell type with motile cilia. Whether these motor ciliated cells are used in locomotion remains unknown.

The general aspect of the external epithelium of *I. pulchra* could be observed with immunostainings against acetylated tubulin which reveals the density of cilia on the external epithelium of the animal (Fig 5C). A commercial antibody directed against the Delta protein from *Drosophila* revealed the morphology of the cells that compose the epithelium and we therefore used it as marker to delineate their shape. Together with phalloidin staining this antibody shows the disposition of actin-layered pores that may be involved in secretion (Fig 5C).

SmFISH for *adhesive plaque matrix protein* (secretory cells) and *SCO-Spondin 3* (Secretory cells) and *Fucolectin-6* (Epithelial cells) seemed to show scattered patterns of expression throughout the superficial layer of the body with slightly higher occurrence in the anterior half of the animal (Fig 5D). This suggests that secretory cells in acoels are not only grouped in specific secretory glands (Klauser, 1986, Pedersen, 1965) but can also be present throughout the epithelium. Additionally, the expression of cadherins and protocadherins in some secretory cells may indicate that they use these proteins to attach to the epithelium. Based on the data collected for different types of secretory cells, it seems probable that, at least, some of these cells are involved in processes of active external secretion (i.e mucus; Klauser 1986) or in innate defense mechanisms.

### Muscle cells show high marker conservation with other bilaterians

Specific genes involved in the formation of contractile fibers enabled the characterization of three clusters of muscle cells (Ladurner & Rieger 2000; Rieger & Ladurner 2003; Raz et al. 2017). Major components of contractile fibers such as *myosin heavy chain, myosin light chain, troponin, sarcalumenin* and *tropomyosin* are broadly expressed in two of those clusters (Fig 6A). Interestingly, certain markers suggest similarities between muscle cells and epithelial cells. For instance, laminin, which is a major component of the basal lamina of epithelia appears here to be broadly expressed in muscle cells. These similarities are reflected on the UMAP plot in which the main muscle cell clusters is relatively close to epithelial cells with the consistent appearance of a smaller cluster that seems to bridge clusters of muscle and epithelial cells (Fig 6B). This latter cluster would suggest the presence of a set of muscle cells that share markers with epithelial cells (*cadherin, protocadherin-1, protocadherin-2 and fucolectin-6)* which are mostly involved in cell-cell adhesion. This could indicate that these muscle cells are anchored to the epithelium through cadherins. Since acoels rely on the sole use of cilia of epithelial cells for locomotion it is likely that there must a close coordination between the function of the ciliated epithelium and contractile muscle to modulate movements.

**Figure 6:**
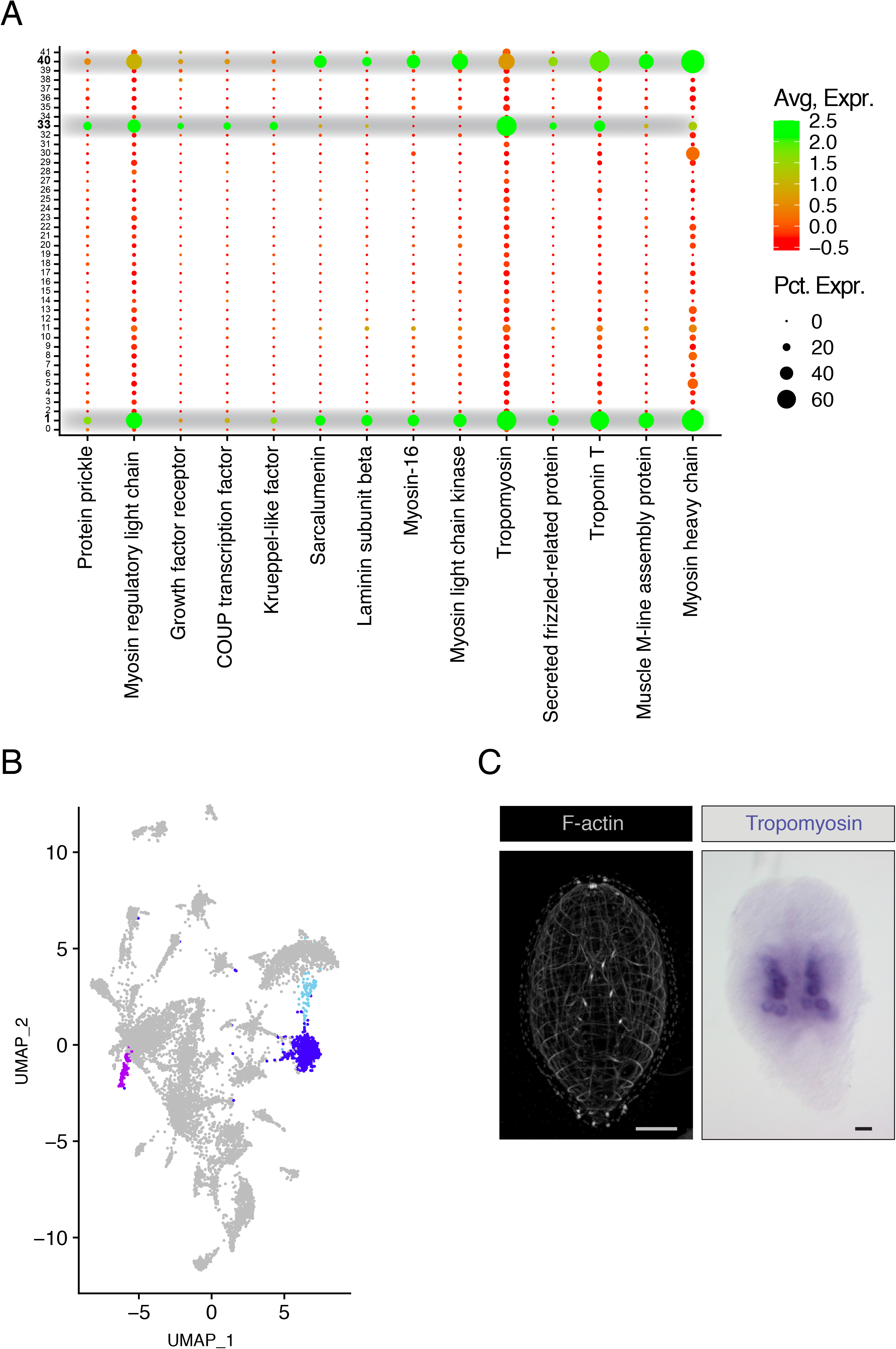
Muscle cells show high marker conservation with other bilaterians. (A) DotPlot showing the expression of specific markers for muscle across all 42 cell clusters. Three likely candidates for muscle cells are highlighted. (B) UMAP highlighting the three clusters identified as muscle cells. (C) Phalloidin staining showing the actin filament network of *I. pulchra* and ISH for tropomyosin. Scale bars = 20μM.

Muscle cells are one of the few cell types in which defining transcription factors can be identified in our dataset: The transcription factors *krueppel-like* and *COUP* are specifically detected in muscle clusters. In addition, the *wnt* interacting partner *frizzled* is specifically expressed in all three of these muscle clusters (Fig 6A). The whole structure the muscle network of *I. pulchra* can be observed with phalloidin staining of actin filaments (Fig 6C). Additional ISH for tropomyosin showed high expression in or around the gonads (Fig 6C).

Interestingly, most of these presumed muscle cells also express several types of acetylcholine receptors (*AChR-4, AChR-5, AChR-6*), indicating that the neuromuscular junctions are likely mediated by cholinergic neurons (see: Fig 3).

### Cell-type markers are conserved within the Xenacoelomorpha

In an attempt to detect the presence of both Acoela-specific and Xenacoelomorpha-specific cell types or signatures, we extracted the sequences of *I. pulchra* for which we could not identify orthologs in other clades and compared them to other Xenaceolomorpha transcriptomes we have produced in the laboratory (*Symsagittiera roscoffensis* and *Xenoturbella bocki*). As a result, we identified 4332 unknown sequences that have homologous sequences in *S. roscoffensis* and 3063 in *X. bocki*. We observed that all Xenacoelomorpha-specific sequences found among the cell-type markers were also present in the group “Acoela-specific”, with only one exception (*DN18593*), suggesting that these genes’ sequences are well conserved among Xenacoelomorpha, even though they do not have similarities with sequences of other phyla. They represent a collection of very derived sequences that could still represent orthologues of other genes but highly modified or alternatively represent Xenacoelomorpha-specific novelties. The fact that they are expressed in all three species also indicates that they are likely to be functionally relevant. We initially looked for these genes in the group of “uncharacterized” cell types but the fraction of Acoela- and Xenacoelomorpa-specific sequences in these clusters was not higher than in other clusters with assigned phenotype. This prompted us to avoid characterizing them as novel Xenacoelomorpha-specific cell types. However, the relevance of these genes should not be discounted since they are abundantly present in many of the better characterized cell types, particularly in digestive, epithelial and secretory cells plus also in some of the sensory cell types.

To conclude, the expression of well-known orthologs of bilaterian cell markers in this dataset has been sufficient to classify cell types into different functional categories. However, and surprisingly, it appears that each of these categories show significant expression of transcripts shared only within Xenacoelomorpha (clade-specific transcripts). At this point, however, we cannot rule out the possibility that *I. pulchra* have some specific cell types, though the data would better fit a model in which all cell types of Acoela show some unique transcriptional profiles different from those of the remaining bilaterians. Further analysis of the genome of *Isodiametra pulchra* as well as other members of the clade could help clarify that issue and more accurately describe gene conservation among Xenacoelomorpha and their putative relationship with those of other phyla (Guijarro-Clarke et al. 2020).

## Discussion

The insights into cell-type diversity through single-cell RNAseq experiments provides a powerful way to approach cell-type evolution in a highly reproducible manner. In this study, we provide a cell atlas of the juvenile acoel *Isodiametra pulchra* that displays functionally distinct cell types. This is the first time that a high-resolution expression atlas is provided for any member of the enigmatic group Xenacoelomorpha. While the amount of predicted cell types is consistent with what has been morphologically characterized in the past, the newly characterized subsets of cells and the specific genes expressed in each of these subsets offer valuable tools to further characterize the histology and developmental trajectories of these animals.

The nervous system of *I. pulchra* could now be described at the level of individual cells providing additional information about the chemical modalities of neurotransmission of acoel neurons. We identified distinct types of sensory cells that could be involved in chemosensation, mechanosensation and possibly photosensation. In addition to improving our understanding of the nervous system of acoels, this provides an entry point to functional and behavioral studies in which the described sensory markers could be stimulated or tampered with to better understand the specific role of each sensory cell type. The enormous plasticity of nervous system architectures in the Xenacoelomorpha has now a cellular reference frame for us to understand the building blocks that give rise to this great diversity.

We showed that digestion and transport of nutrients was probably carried out by a variety of cell types that secrete different digestive enzymes, including several members of the cathepsin family. These digestion processes are likely to act together with the digestive syncytium, though it remains unclear whether the enzymes are secreted into the syncytium of act as processes of intracellular digestion. This can be the case with certain types of cathepsins that are predominantly present in lysosomes in various species (Kirschke et al. 1995). The detection of several digestive enzymes opens the possibility of better understanding one of the most enigmatic tissues of Xenacoelomorpha, the gut (Gavilán et al. 2019). Other cell types were shown to express high levels of nutrient and ion-transport related transcripts, indicating a possible function in nutrient absorption and active distribution of these nutrients to other cells. In addition, one cell cluster expresses markers that have been previously proposed as having a role in excretory processes in *I. pulchra* (Andrikou et al., 2019), suggesting an involvement of these cells in the elimination of metabolic waste. Interestingly, some cell types simultaneously express markers of both digestive enzyme and nutrient/ion transport. This could indicate that these cells have dual roles (Gazizova et al. 2017) or that we have captured progenitors that later on give rise to two phenotypically distinct cell types. Since very little is known on the maturation process of the gut, both alternatives remain possible. Together, these results show for the first time that the digestive system of acoels consists of more than just a simple, homogeneous, digestive syncytium but that it encompasses many cell types that seem to assume different and distinct roles for digestion.

We provide new information about the epithelia of *I. pulchra* by describing two main categories of epithelial cells whose phenotypes suggest distinct functions. One category seems to express more classic structural elements of epithelia while the other express markers of motor cilia which suggests an involvement in the locomotory behavior, which in acoels relies exclusively on ciliary motion. Our data suggests that the movements of these motor cilia are mediated by axonemal dyneins known to be a major component of cellular motors in many eukaryotes including unicellular protists (King 2012).

The presence of secretory cells and glands in different Xenacoelomorpha have been thoroughly described at the morphological level (Pedersen 1965; Klauser 1986), with mucus secretion in *Symsagittifera roscoffensis* (Acoela) proposed as an aid to locomotion (Martin 2005). In this study we extended this knowledge by describing the diversity of secretory cell types that suggest a broad variety of functions such as adhesion and defense against predators or pathogens. This first molecular description of secretory systems of an acoel provides important elements to compare secreted products such as bioadhesive proteins and toxins to those of other animals and follow their expression over evolutionary time (Tyler 1976).

The peculiarity of acoel cell types compared to other bilaterians could be anticipated given the long time that the clade has evolved independently and the fast rate of nucleotide substitution that characterize their genomes. This is a group an ancient bilaterians that diversified around 500-600 Mya. This combination of old clade diversification and the fast rate evolution of their genomes may obscure the similarity of their genes with those of other animals, resulting in an added difficulty for detecting sequence similarities. However, and in spite of the contribution of these factors, our results show that instead of predicting a large number of novel cell types, *I. pulchra* displays an array of known cell types that express a combination of some conserved markers plus some clade-specific ones. Many of the latter sequences are, indeed, conserved across the different Xenacoelomorpha pointing to the presence of some specific functions carried out by conserved cell types. This shows that despite their important diversity, Xenacoelomorpha possess some clade-specific conserved genes as is the case for other phyla (Paps and Holland 2018). How these sequences contribute to the specific character of the Xenacoelomorpha cell types remain to be studied.

Together, our results pave the way for the further analysis of the different roles that these different cell types have in the morphology and physiology of acoels. Moreover, our results identified candidate markers for elusive cell populations such as multipotent stem cells or secretory cells and help us build a molecular map of most organ systems. For the first time, we provide a thorough analysis of gene expression in acoel individual cells. The description of a catalog of cell-types in *Isodiametra pulchra* should help us understand bilaterian evolution through the perspective of its cellular constituents. This is a powerful way to access to the constructional principles that guide the different morphologies of Xenacoelomorpha or any other animal, by helping to understand the diversity and arrangement of their cellular building blocks. The presence of cell types with mixed signatures and the generalized usage of clade specific transcripts are especially relevant since they should explain the specificities of Xenacoelomorpha tissues and their functional activities. With the data provided here and the implementation of cross-species analysis of single cell transcriptomic data, the possibility of tracing the evolutionary histories of cell types becomes a reachable objective. Single-cell data and its translation into cell type characters, will be of special interest for also tracing the evolutionary history of many clades. We hypothesize that this source of data should help us, in addition, to understand the phylogenetic affinities of Xenacoelomorpha.

## Materials and methods

### Animal culture and breeding

Animals were kept at 20°C in glass petri dishes. They were fed by being transferred on a freshly grown biofilm of the diatom *Nitzschia curvilineata* every 6 weeks. Hatchlings were collected and identified for the experiment based on size. All animals were starved few hours prior to RNAseq experiments to avoid excessive algae contamination.

### Single-cell suspension

Whole animals (~100 hatchlings) were dissociated by incubating for 1h at 25°C in a collagenase solution (1mg/mL, Sigma-Aldrich C9722) with continuous agitation. The suspension bas briefly vortexed every 10min to ensure full digestion. The suspension was briefly centrifuged (5min, 750rcf) and the pellet was suspended in RNAse-free PBS with 0.04% BSA (Thermo Fischer Scientific AM2616). Cells were filtered through a 40μM Flowmi Cell strainer (Bel-Art H13680-0040). Centrifugation and resuspension in PBS were repeated to wash the cells. The whole suspension was then gently pipetted up and down around 200 times coated pipette tips. The cell concentration was estimated using a hemocytometer (Neubauer improved – Optik Labor) under a binocular microscope.

### Transcriptome assembly and annotation

RNA was extracted from animals in mixed stages, RNA was purified using a Qiagen RNA purification kit. The poly-adenylated transcriptomes (mRNA) of I. pulchra, S. roscoffensis and X. bocki were sequences on an Illumina HiSeq 3000. We generated a total of 84 930 312, 61 857 037, and 116 117 207 paired-end 150 bp long reads for pulchra, S. roscoffensis, and X. bocki, respectively, and these data sets have been uploaded to NCBI (accession numbers for Ip: SAMN07276911, Sr: SAMN07276888, and Xb: SAMN07276887). De Novo assembly was performed using the TRINITY pipeline and annotation was done using TRINOTATE (v3.1.1, Bryant et al., 2016). BLAST (v2.7.1) homology pairing was performed against the SwissProt database. The name assigned to genes in this study corresponds to the Blastx result with lowest e-value. Protein domain (Pfam) search was done using HMMER (v3.1b2).

The poly-adenylated transcriptome of *Isodiametra pulchra* was generated on an Illumina HiSeq 3000 and generated 84,930,312 paired-end 150bp long reads (Brauchle et al., 2018). This initial transcriptome assembly of *Isodiametra pulchra* revealed ~300,000 transcripts. The very high number of identified transcripts was determined to be caused by both the presence of shorter reads that could not be confidently mapped to other transcripts and because of the existence of many transcriptional isoforms for certain genes. To enable better mapping of the reads obtained in single-cell RNA sequencing experiments we reduced redundancy in our transcriptome . The final number of transcripts in the non-redundant transcriptome is of 45,000.

Completeness of the transcriptome was tested by BLASTing the transcriptome against BUSCOs (Benchmarking Universal Single Copy Orthologs, Simão et al., 2015, Seppey et al., 2019, v3.0.2) as defined for all Metazoa (978 genes). Out of these 978 genes, 732 were found in our non-redundant transcriptome, either as a single-copy or as duplicated genes, 61 were fragmented and 185 were not found (Supp. fig S1B). This corresponds to an estimation of 74.8% of BUSCO groups that could be identified in our non-redundant transcriptome, either as a single-copy or as duplicated genes. The list of missing BUSCOs is available in supplementary table 3. We performed the same analysis our redundant transcriptome and obtained better results with 326 single-copy, 542 duplicated BUSCOs, 25 fragmented and 85 missing. This showed a higher proportion of identified BUSCOs (88.7%) but with a very high proportion of duplicated genes (55.4%), indicating the important redundancy of this transcriptome.

Functional annotation based on Blastx similarity searches in a general protein database (Swissprot) and a database of curated protein domains (Pfam) was performed on the assembled transcriptome. About half of these transcripts had clear similarities to proteins of the database (e-value ≤ 0.01) and/or a predicted protein domain. The annotation reveals that the *I.pulchra* transcriptome contains 20,446 sequences with orthologous in other organisms leaving 24,554 transcripts of unknown identity (Supp. fig S1C). 17,810 transcripts encode for a protein (Open reading frame) with a conserved structural domain. The others might correspond to acoel-specific or highly divergent sequences. To assess the conservation of these unknown genes/sequences within the Xenacoelomorpha and verify that they are not contamination of our transcriptome, we generated transcriptome assemblies for *Symsagittifera roscoffensis* (Acoelomorpha) and *Xenoturbella bocki* (Xenoturbellida) and blasted the unknwown sequences from *I. pulchra* (e-value ≤ 0.01) against them This process identified 4332 transcripts that have orthologues with *S. roscoffensis* and therefore may be acoel-specific and 3063 transcripts that have orthologues in *X. bocki* and might therefore be Xenacoelomorpha-specific (Supp. fig S1A, supp. tables 4&5).

### 10x genomics

Single-cell RNA sequencing experiments were all performed at the Next Generation Sequencing platform of the university of Bern. scRNA-seq libraries were prepared using the Chromium Single Cell 3’ Library & Gel Bead Kit v2 or v3 (10X Genomics), according to the manufacturer’s protocol (User Guide). Chips were loaded after calculating the accurate volumes using the “Cell Suspension Volume Calculator Table”. With an initial single-cell suspension concentration estimated at 300 cells/μl, we targeted to recover approximately 8000 cells. Once GEMs were obtained, reverse transcription and cDNA amplification steps were performed.

Sequencing was done on Illumina NovaSeq 6000 S2 flow cell generating paired-end reads. Different sequencing cycles were performed for the different reads, R1 and R2. R1, contained 10X barcodes and UMIs, in addition to an Illumina i7 index and R2 contained the transcript-specific sequences. The total of reads for the combined experiments was 734,346,794 post-normalization resulting in an average of 49,404 reads per cell.

ScRNA-Seq Cell counting and viability assessments were conducted using a DeNovix CellDrop Automated Cell Counter with an Acridine Orange (AO) / Propidium Iodide (PI) assay. Thereafter, GEM generation & barcoding, reverse transcription, cDNA amplification and 3’ Gene Expression library generation steps were all performed according to the Chromium Single Cell 3’ Reagent Kits v3 user Guide (10x Genomics CG000183 Rev B). Specifically, 37.3 µL of each cell suspensions (300 cells/µL) were used for a targeted cell recovery of 7000 cells. GEM generation was followed by a GEM-reverse transcription incubation, a clean-up step and 12 cycles of cDNA amplification. The resulting cDNA was evaluated for quantity and quality using fluorometry and capillary electrophoresis, respectively. The cDNA libraries were pooled and sequenced paired-end and single indexed on an illumina NovaSeq 6000 sequencer with a shared NovaSeq 6000 S2 Reagent Kit (100 cycles). The read-set up was as follows: read 1: 28 cycles, i7 index: 8 cycles, i5: 0 cycles and read 2: 91 cycles. An average of 521,045,087 reads/library were obtained, equating to an average of 65,130 reads/cell. All steps were performed at the Next Generation Sequencing Platform, University of Bern.

### 10x data processing

The single-cell sequencing data was processed with Cell Ranger (10x genomics, v3.0.2) using the provided pipeline. The reference sequence was obtained by concatenating the transcriptome in one single sequence with each transcripts separated by 1000 Ns. Two separate experiments using different BeadKits were batched together using the provided aggregation function. This resulted in a total of 14,864 cells with a median of 405 genes/cell for a total of ~20,000 genes in total.

### Seurat data processing

Seurat pipeline (v3.1.4, Satija et al., 2015, Butler et al. 2018) was adapted and used on our dataset. Cells were filtered to include only those with a gene per cell count of 200 to 2000. Seurat objects were log-normalized with a scale factor of 10,000. Variable genes were identified with the FindVariableGenes function with low and high X cutoffs of 0.0125 and 3 respectively and a Y cutoff of 0.5. The dataset was then scaled by applying linear transformation. 25 principal components were selected for clustering based on an ElbowPlot test. Clustering was performed with a PCA reduction with a resolution of 2.5. UMAP dimensional reduction was used to plot clusters.

### Immunostainings

Whole animals were fixed for 30min in 4% formaldehyde. Primary antibodies were used in the following concentrations (mouse anti-dSAP47 1:20, rabbit anti-5HT 1:200, mouse anti-delta 1:200, mouse anti-acetylated tubulin 1:500). Primary antibodies were incubated at 4°C overnight and secondary antibodies were added for 2 hours at room temperature (concentration 1:200). Images were taken with a Leica SP5 confocal microscope.

### smFISH

Probes were synthetized by Stellaris® (Biosearch technologies). Animals were fixed for at least 1 hour in 4% formaldehyde at room temperature and subsequently washed with PBS and transferred in 100% methanol to be stored at −20°C. Specimens were rehydrated in 1:3, 2:3 and 1:1 PBT. In situ hybridization was done following the provided protocol for drosophila embryos (*Orjalo et al., 2016*) using hybridization temperature of 45°C instead of 37°C and reducing the reaction volumes to 100uL per probe mix. Images were taken with a Leica SP5 confocal microscope.

### InSitu hybridization

Animals of all stages were collected from the dish and put in an Eppendorf tube. They were anaesthetized with 7 % MgCl and fixed with 3.7 % paraformaldehyde in PBTx (0.3 % Triton X-100 in 1x PBS) for 30 min at room temperature. The washes were done 3 x with PBTx and then dehydrated in methanol. The samples were stored at −20°C for up to several months.

Samples were rehydrated in Eppendorf tubes in four steps from 100% methanol to 100% PBT (0.1 % Tween-20 in 1x PBS) and washed 3 × 5min in PBT. A proteinase K treatment was applied for 8 min at a concentration of 20 µg/ml in PBT. The proteinase K activity was stopped with 2×5min washes in glycine (4 mg/ml in PBT, *Roth*). A series of 5 min washes were performed on a shaker set to 30 rpm in the following order: 1% triethanolamine in PBT, 0.25 % acetic anhydride in 1 % TEA, 0.5 % AA in 1% TEA followed by three washes in PBT. Samples were refixed in 3.7% paraformaldehyde solution was used for 20 min at room temperature followed by 5 washes of 5 min in PBT.

Before the pre-hybridisation the samples were placed in a 50/50 solution of hybridisation buffer and PBT for 10 min at RT. The recipe for the hybe buffer was the following : 50% deionised formamide, 5x SSC (0.75M sodium chloride + 0.075M sodium citrate), 1% SDS, 0.1% Tween-20, 50 µg/ml heparin, 100 µg/ml herring sperm, 0.01M citric acid. Afterwards the samples were kept in 100% hybe buffer for 2h at 60°C. The probes were added after having been denatured at 90°C for 5 min and added to the flask to a final concentration of 1000ng/ml. The hybridisation was performed at 60°C in a water bath for two days.

The first steps after the hybridisation were done at 60°C in a water bath and in the following solutions: 100% hybe buffer, 75% hybe buffer 25% 2X SSC, 50% hybe buffer 50% 2X SSC, 25% hybe buffer 75% 2X SSC 5 min each and then 2 × 30 min in 2X SSCT. The samples were blocked in 1x maleic acid buffer blocking solution for 3h at 4°C. The antibody incubation was performed with a 1: 2000 anti-DIG-AP (Roche) in blocking solution for 12h at 4°C. For staining, an NTMT solution was used with 1:100 NBT/BCiP. Primers used in this study were the following:

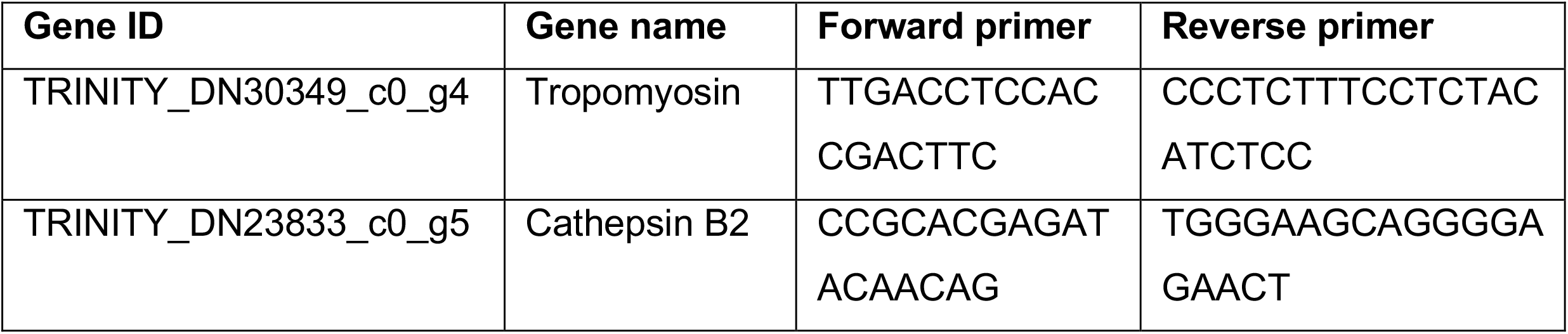

## Acknowledgements and funding information

We thank Dr. Pamela Nicholson and her team at the next generation sequencing facility (NGS) of the university of Bern for their expertise and their help with the single-cell RNA transcriptomics experiments.

This work was supported by the Swiss National Science Foundation to SGS (grants IZCOZ0_182957 and 310030_188471) and by the Agencia Estatal de Investigación to PM (grant PGC2018-094173-B-I00).

## Data availability

Raw data and processed datasets for single-cell transcriptomics are available on NCBI (accession number GSE154049).

## Tables

All tables are available as supplementary data.

**Supplementary table 1:** List of all genes mentioned in the text with their corresponding name as found in the transcriptome as well as the best BLASTx hit from Swissprot.

**Supplementary table 2:** List of the top 10 markers in each cluster and the corresponding gene name and BLASTx hit.

**Supplementary table 3:** List of all 978 BUSCOs used to assess our non-redundant transcriptome and the corresponding *Isodiametra pulchra* transcript if present.

**Supplementary table 4:** List of all uncharacterized *Isodiametra pulchra* genes with similar sequences in *Symsagittifera roscoffensis*.

**Supplementary table 5:** List of all uncharacterized *Isodiametra pulchra* genes with similar sequences in *Xenoturbella bocki*.

---

**Supplementary figure 1:**
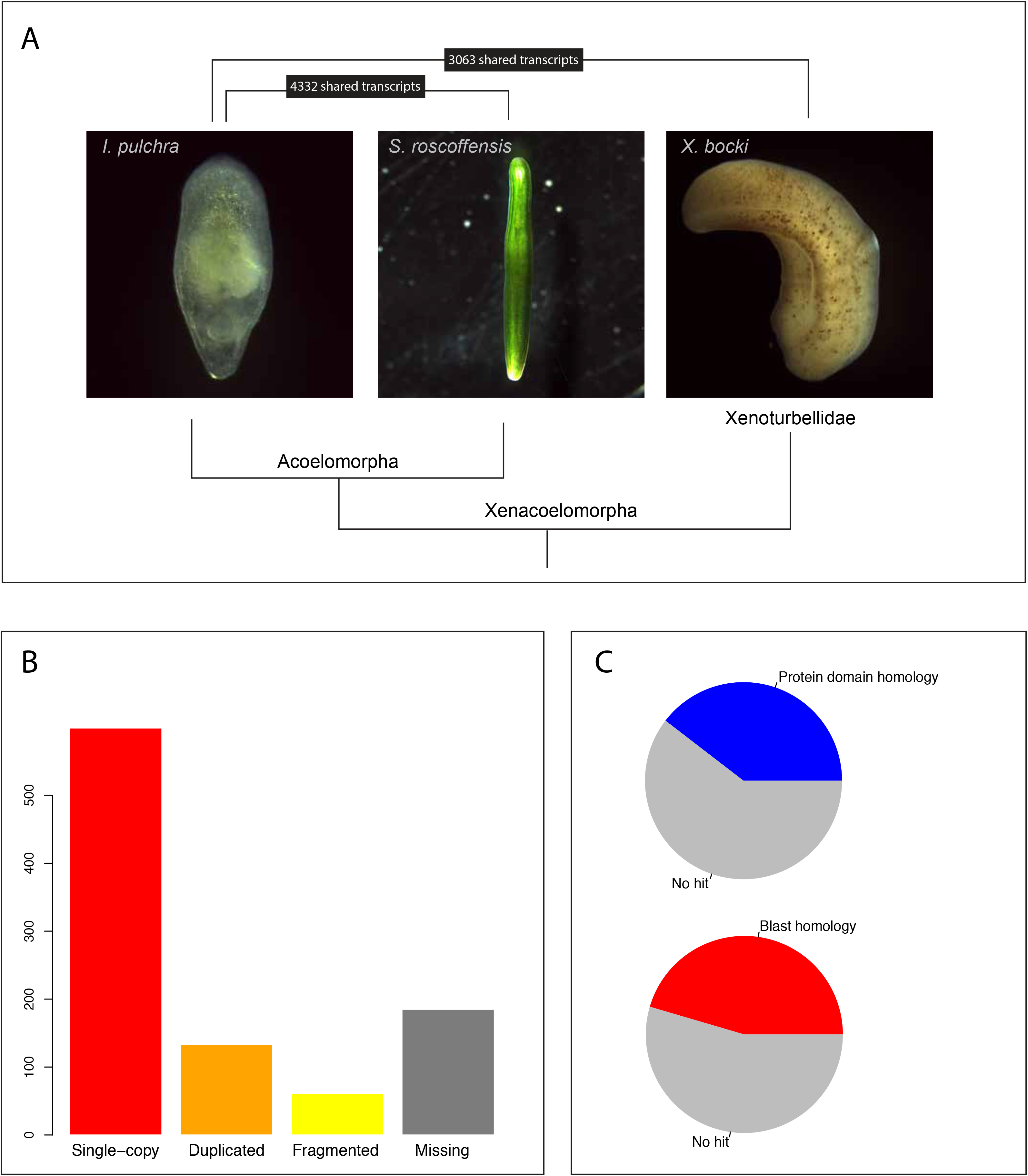
Transcriptome annotation and comparisons. (A) Photographs showing three representatives of Xenacoelomorpha: *Isodiametra pulchra*, *Symsagittifera roscoffensis* and *Xenoturbella bocki.* Their phylogenetic relashionship is shown underneath and the number of identified clade-specific candidate transcripts with *I.pulchra* are shown above. (B) Report of alignements of the non-redundant *Isodiametra pulchra* transcriptome to the Benchmark Universal Single-copy Orthologs (BUSCOs) of the metazoan dataset of 978 genes. (C) Pie plots indicating the proportion of the transcriptome with either Protein domain homology or BLASTx-based homology in Swissprot database.

**Supplementary figure 2:**
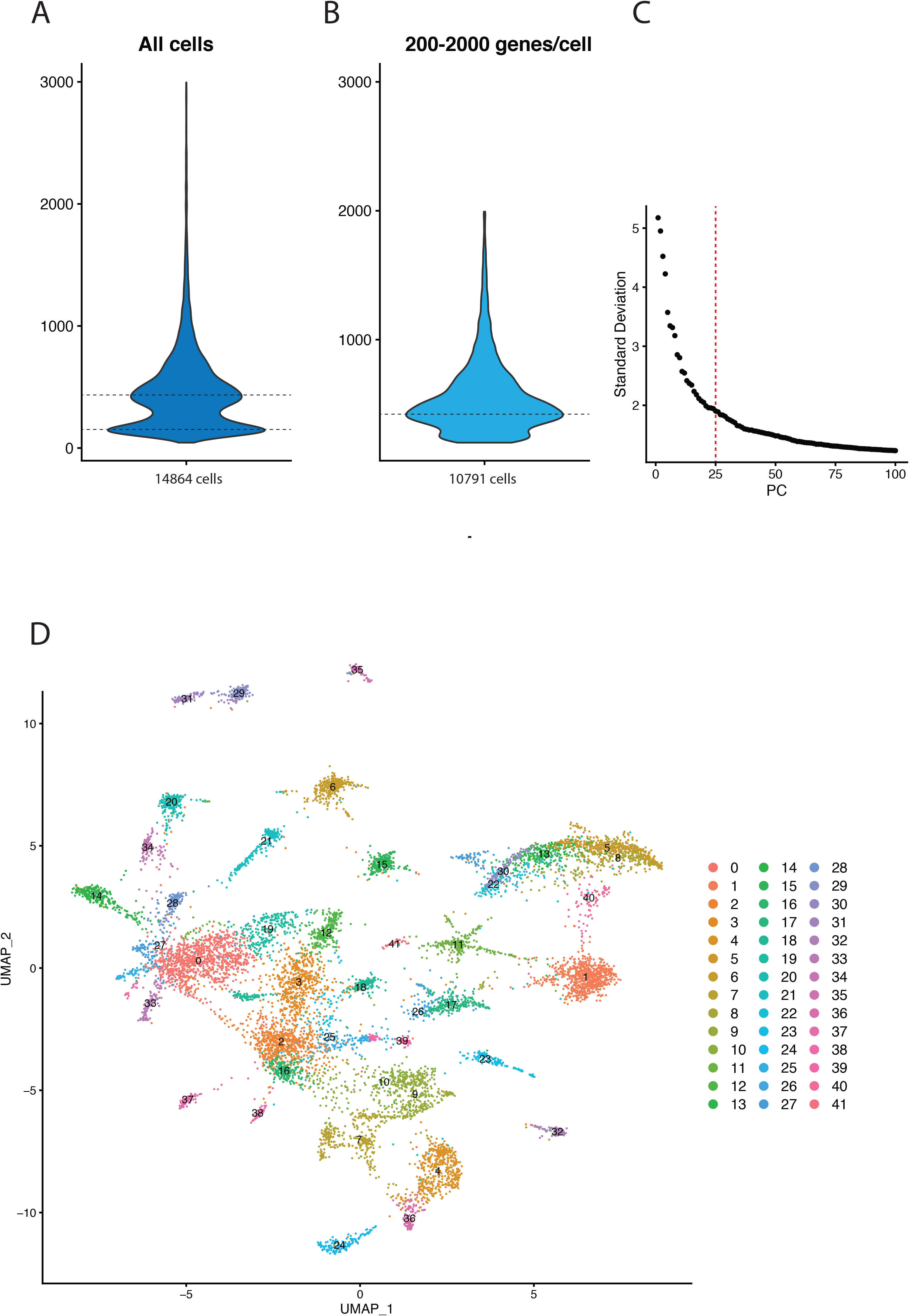
Quality controls and raw clusters. (A) Violin plot showing the distribution of all cells based on their gene-per-cell counts. (B) Violin plot showing the distribution of filtered cells cells based on their gene-per-cell counts. Only cells with 200 to 2000 genes are kept to improve overall cell quality homogeneity. (C) Eblow plot showing the raking of principal components (PCs) based on their standard deviation. Red dotted line shows the upper limit of 25 PCs used for further analysis, corresponding to the elbow of the curve. (D) UMAP of the raw 42 clusters obtained on Seurat by using 25 PCs and a resolution of 2.5.

**Supplementary figure 3:**
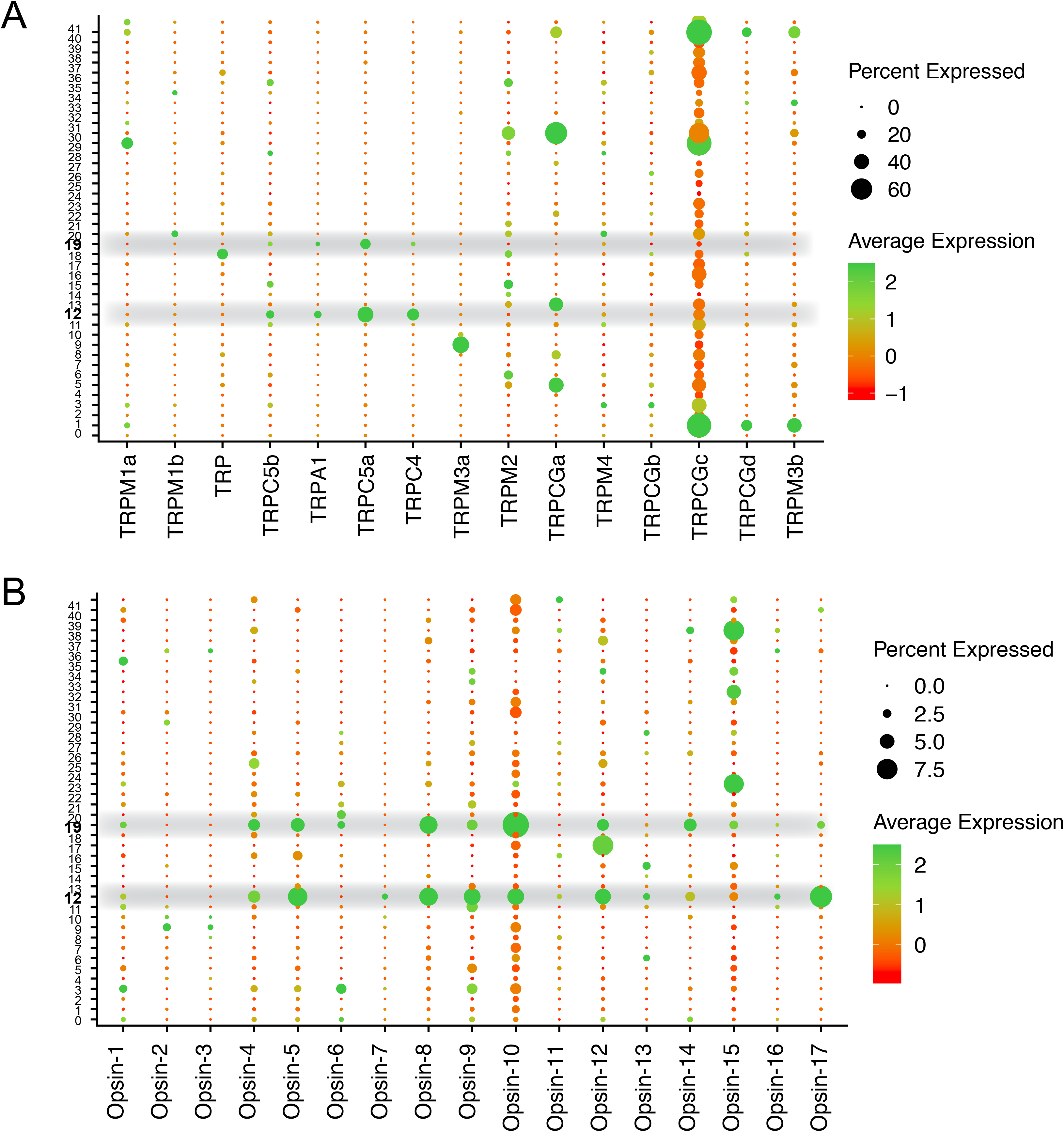
TRP channels and Opsins. **(A)** Dotplot showing the expression of all identified TRP channels across all clusters. Highlighted clusters are identified as TRP^+^ neurons. **(B)** Dotplot showing the expression of all identified opsins across all clusters. Highlighted clusters are identified as TRP^+^ neurons.

## Notes

### Competing Interest Statement

The authors have declared no competing interest.

